# Conditional U1 Gene Silencing in *Toxoplasma gondii*

**DOI:** 10.1101/008649

**Authors:** Manuela S. Pieperhoff, Gurman S. Pall, Elena Jiménez-Ruiz, Sujaan Das, Eleanor H Wong, Joanne Heng, Sylke Müller, Michael J Blackman, Markus Meissner

## Abstract

In absence of powerful siRNA approaches, the functional characterisation of essential genes in apicomplexan parasites, such as *Toxoplasma gondii* or *Plasmodium falciparum,* relies on conditional mutagenesis systems. Here we present a novel strategy based on U1 snRNP-mediated gene silencing. U1 snRNP is critical in pre-mRNA splicing by defining the exonintron boundaries. When a U1 recognition site is placed into the 3’-terminal exon or adjacent to the termination codon, pre-mRNA is cleaved at the 3’-end and degraded, leading to an efficient knockdown of the gene of interest (GOI). Here we describe a simple one-step approach that combines endogenous tagging with DiCre-mediated positioning of U1 recognition sites adjacent to the termination codon of the GOI which leads to a conditional knockdown of the GOI in *Ku80* knockout and RH *T. gondii* tachyzoites. Specific knockdown mutants of the reporter gene GFP and several endogenous genes of *T. gondii* including the clathrin heavy chain gene 1 *(chc1)*, the vacuolar protein sorting gene 26 (*vps26)*, and the dynamin-related protein C gene (*drpC)* were silenced using this new approach. This new gene silencing tool kit allows protein tracking and functional studies simultaneously.

## Introduction

Protozoan parasites of the phylum Apicomplexa, such as *Toxoplasma gondii, Plasmodium spp.* or *Cryptosporidium spp.*, are important pathogens of human and livestock populations. These organisms also represent fascinating biological systems with unique organelles and adaptations to an intracellular lifestyle. With completion of genome sequencing for several apicomplexans (www.eupathDB.org) and combined efforts to characterise the transcriptome, proteome and metabolome, the list of promising drug and vaccine candidates is increasing. However high-throughput analyses of genes of interest (GOI) are lacking since current reverse genetic technologies are time consuming [1].

While conditional mutagenesis is feasible in *T. gondii* and *P. falciparum* using several key technologies, such as the Tet-inducible [2,3], protein destabilisation [4,5] or DiCre-recombination systems [6,7], the generation of conditional mutants for essential genes needs to be more streamlined and standardised to enable a higher throughput. Therefore, time-consuming obstacles, such as cloning complex knockout (KO) constructs and using strategies that involve the generation of multiple transgenic parasite lines in order to localise and knockdown a single protein of interest need to be overcome.

Here we describe a novel technology that combines the advantages of C-terminal tagging of a GOI in a *T. gondii Ku80* background [8,9] with the DiCre-dependent placement of U1-recognition sites at the terminal exon of the GOI and therefore efficient knockdown of the GOI’s transcripts.

Previous studies in human cell lines have established the critical role of U1 small nuclear ribonucleic particles (snRNP) in splicing of pre-mRNA. In particular by defining the 5’ donor site of an intron [10], it has also been demonstrated that it can block accumulation of a specific RNA transcript when it binds to a donor sequence within the terminal exon (i.e. close to the STOP codon) of that GOI [11]. Earlier studies demonstrated the feasibility of achieving specific and tight regulation of expression levels either by directing modified U1 snRNA to a unique sequence within the terminal exon [12] or by designing GOI-specific U1 adaptors that target the terminal exon of a GOI sequence and contain the U1 domain, thereby leading to recruitment of the U1 machinery resulting in pre-mRNA degradation [13]. While the former technology requires multiple genetic manipulations (positioning of U1 recognition sequences into the terminal exon of the GOI or expression of modified U1 snRNA [12,14]), U1-adaptors can be used as a promising alternative to RNA interference (RNAi) by directly introducing them into the cell [13]. However, our efforts to adapt the synthetic adaptor strategy to *T. gondii* have not yet been successful.

Here we present an alternative to U1 adaptors that can be easily translated to other eukaryotes. We combined the high efficiency of regulated DiCre-mediated recombination [6,7,15] with endogenous tagging and U1 mediated knockdown of target gene expression. We used the apicomplexan parasite *T. gondii* as a model system to demonstrate the feasibility of this approach. This novel technology has several advantages, including easy vector design, a requirement for only a single genetic manipulation for both localisation and knockdown studies, and high efficiency of specific gene knockdown. We also tested this system in the related apicomplexan parasite *P. falciparum*. Surprisingly, a direct adaptation of the strategy failed in this parasite, suggesting the existence of crucial differences in mRNA splicing mechanisms between these two genera.

## Results

### Positioning of U1 recognition sites within the terminal exon of a GOI

Since the spliceosome and the mechanisms involved in the definition of exon-intron boundaries are highly conserved in eukaryotes [10], we reasoned that positioning U1 recognition sequences in the terminal exon of a *T. gondii* GOI would result in efficient gene knockdown, as previously shown in other eukaryotes [31]. We first compared the 5’-end of U1 snRNAs of apicomplexan parasites with other eukaryotes, since the first 10-nt of U1-snRNA recognise the 5’-splice site (Figure 1A and 1B). We confirmed that the recognition sequence is highly conserved and only a single nucleotide substitution at position 2 was identified in *T. gondii* and *P. falciparum* (Figure 1A). We therefore speculated that positioning of a slightly modified 5’-recognition sequence (CAG/GTAA**GTT** instead of CAG/GTAAGTA) should lead to a block in polyadenylation and consequent degradation of the pre-mRNA, resulting in an effective knockdown of expression levels of a GOI (Figure 1B).

**Figure 1.**
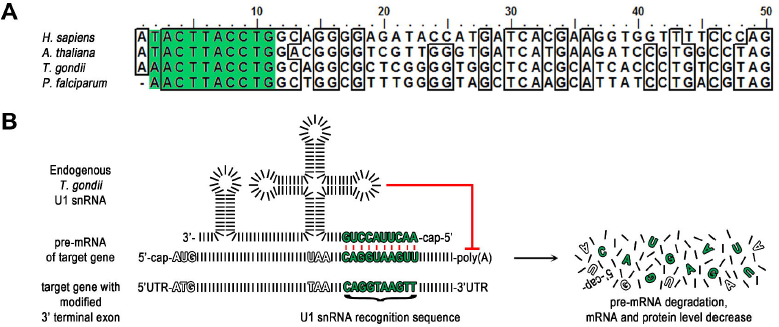
U1 gene silencing strategy in *T. gondii.* (A) Alignment of the first 50 nucleotides of the U1 snRNAs of the indicated organisms. Shaded in green are nucleotides 2-11 complementary to the artificial U1 snRNA recognition sequence in (B). Nucleotides 3-11 are identical across the aligned species. *H. sapiens, Homo sapiens; A. thaliana, Arabidopsis thaliana; T. gondii, Toxoplasma gondii; P. falciparum, Plasmodium falciparum.* (B) Schematic of strategy. Modification of the target gene’s 3’ terminal exon by insertion of a U1 snRNA recognition sequence leads to recruitment of the U1 snRNP to the pre-mRNA of the target gene. Base pairing between these 10 complementary nucleotides of the U1 site and the U1 snRNA of the U1 snRNP occurs and the resulting U1 snRNP pre-mRNA complex inhibits polyadenylation and therefore pre-mRNA maturation. For simplification nucleotides are illustrated as black bars and the target gene is intron-less. UTR, untranslated region.

To test whether this method can be applied in *T. gondii*, based on application experience in mammalian systems with multiple binding sites increasing the level of inhibition in a synergistic fashion [32] two U1 recognition sequences were positioned in tandem directly downstream of the STOP codon of the reporter gene mycGFP (mycGFP-U1, Figure 2A). As a control we also generated an expression vector containing similarly positioned mutant U1 recognition sequences (mycGFP-mutU1, Figure 2A). When these expression vectors were stably introduced into the genome of *T. gondii* RH, GFP was detected when mutated U1 sites were positioned downstream of mycGFP. In contrast, greatly reduced expression of the reporter protein was observed when using the wild type U1 sites positioned downstream of the mycGFP, as shown by immunofluorescence and western blot assays (Figure 2B and 2C). To add temporal control to U1 mediated silencing we took advantage of the DiCre-recombination system, that allows efficient site specific recombination between two loxP sites [7]. We integrated a floxed 3’-UTR of the surface antigen SAG1 followed by two U1 recognition sequences downstream of the mycGFP sequence (mycGFP-floxU1, Figure 2A). Cre mediated recombination was expected to lead to excision of the 3’-UTR and positioning of the U1 recognition sites immediately downstream of the STOP codon of mycGFP. Stable transfection of this construct into RH BloxP DiCre [7] resulted in the generation of green fluorescent parasites, similar to the control population that was transfected with mycGFP-mutU1 (Figure 2B), although immunoblot analysis indicated that introduction of loxP into the 3’-UTR led to lower expression levels of mycGFP (Figure 2C). Upon induction of DiCre activity with 50 nM rapamycin, we observed that almost 100% of all parasites became negative for mycGFP, as determined by immune fluorescence and western blot analysis (Figure 2B, 2C, 2D).

**Figure 2.**
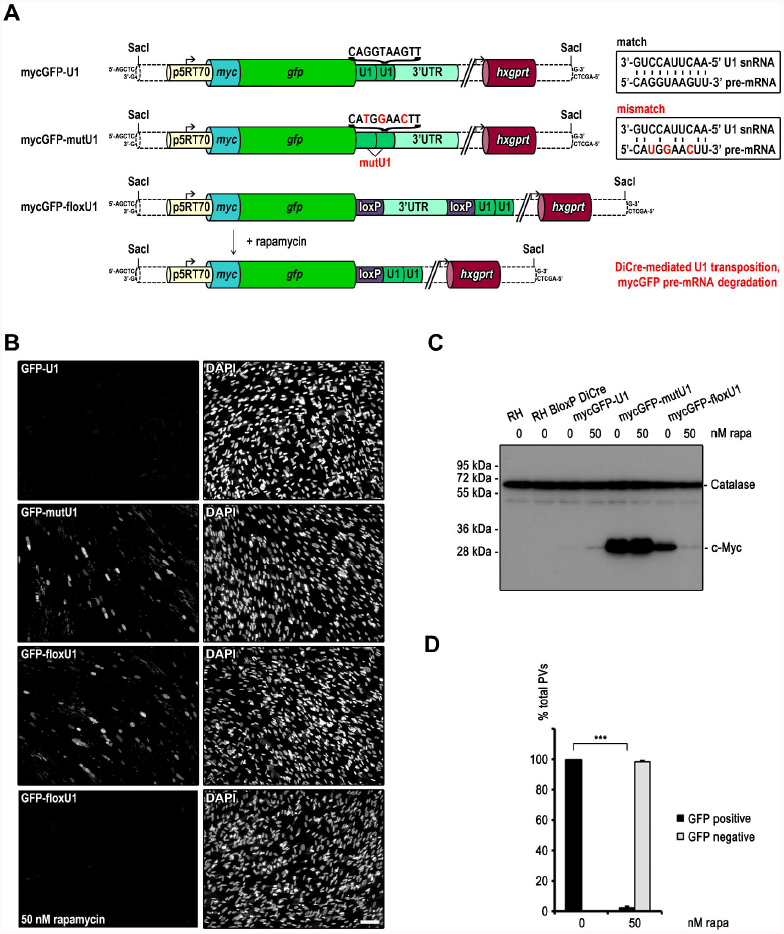
U1 mediated inhibition of reporter gene expression in *T. gondii.* (A) Schematic of reporter gene expression constructs engineered for U1 knockdown strategy. In all three constructs mycGFP expression is driven by the constitutive p5RT70 promoter. In mycGFP-U1 and mycGFP-mutU1 two U1 sites were placed between the STOP codon and the 3’ UTR. In contrast to the wild type U1 site the mutated U1 site (mutU1) confers to gene expression. Respective U1 sequences are indicated above the brackets and matching or mismatching with U1 snRNA are illustrated in the box with mutated nucleotides shown in red. In mycGFP-floxU1 the two U1 sites are placed upstream of the loxP flanked 3’ UTR. Upon addition of 50 nM rapamycin DiCre expression and henceforward site specific recombination is induced. Excision of the floxed linker leads to U1 transposition and reduction of mature mycGFP mRNA levels. For fluorescence (B) and immunoblot analysis (C) of RH BloxP DiCre parasites were stable transfected with the expression constructs shown in (A). Clonal parasite lines were cultivated for 48 hr in absence and presence of 50 nM rapamycin prior analyses. (B) Green, GFP; blue, DAPI; scale bar represents 100 μM. GFP expression is only detectable in mycGFP-mutU1 and mycGFP-floxU1 parasites in absence of rapamycin. (C) Immunoblot probed with anti-c-Myc and anti-Catalase antibodies. Catalase was used as loading control. Only mycGFP-floxU1 parasites show rapamycin dependent mycGFP expression. Note the very faint band obtained for mycGFP-floxU1 under rapamycin conditions is similar in intensity to the ones obtained for mycGFP-U1 with or without rapamycin treatment. (D) Quantification of GFP silencing efficiency. The graph shows the percentage of GFP positive and negative parasitophorous vacuoles (PVs) determined by examination of 200 vacuoles per condition based on fluorescence analyses of mycGFP expressing clonal mycGFP-floxU1 parasites grown for 48 h in absence or presence of 50 nM rapamycin. Values are means ±SD (n=3). Under rapamycin conditions a significant GFP downregulation of 97.5 ± 1.6% was obtained (***, p<0.001, unpaired two-tailed Student’s T-test).

In summary, our results indicate that U1 mediated gene silencing is very efficient in *T. gondii* and can be tightly controlled by DiCre mediated transposition of U1 recognition sites next to the STOP codon of a reporter gene.

### U1 mediated silencing of the clathrin heavy chain (chc1)

Next we tested whether this knockdown approach can be applied for characterisation of essential genes in *T. gondii.* We selected *chc1* as our gene of interest, since we have previously characterised the function of clathrin using a dominant negative approach [33] and hence could compare the phenotypes to exclude off-target effects. We generated a construct that tags the C-terminal end of *chc1* with a *HA-flag* epitope sequence, followed by the floxed SAG1 3’-UTR and the selectable marker, *hxgprt,* followed by four U1 sites (Figure 3A) Transfection of this construct into Δ*ku80::diCre* parasites [7] results in C-terminal tagging of endogenous *chc1*. Upon addition of 50 nM rapamycin, Cre mediated recombination is activated resulting in excision of the selectable marker and positioning of the four U1 sites directly downstream of the stop codon, which should lead to mRNA degradation and knockdown of *chc1* (Figure 3A). We confirmed correct integration of the CHC1 HA-FLAG tagging vector by diagnostic PCR analysis of genomic DNA ([33]; Figure 3A and 3B). Importantly, analytical PCR on genomic DNA isolated from parasites that were treated with 50 nM rapamycin for 24 hours confirmed deletion of the floxed DNA spacer (3’-UTR SAG1 and *hxgprt*). However, excision appears not to be highly efficient, since the non-excised locus was still detectable (Figure 3C). Furthermore, when equal amounts of parasites were analysed in immunoblot analysis 24 hours after rapamycin induction, no differences in protein levels were evident (Figure 3D). Next, a growth analysis was performed, in which induced and non-induced parasites were grown on HFF monolayers for 7 days. No differences in plaque size were apparent (Figure 3E). However, immunofluorescence analysis confirmed that approximately 25% of the parasites showed a significant reduction in CHC1-staining and two distinct parasite populations were readily detected in the induced sample as early as 24 hours after induction (Figure 3G). While ∼75% of parasites showed the typical staining pattern for CHC1-HA, apical to the nucleus in the Golgi area, (see also [33]; Figure 3F), ∼25% of parasites revealed only a weak and diffuse staining pattern, clearly indicating knockdown of *chc1.* This efficiency of target gene excision corresponds well with earlier reports, where *T.gondii ku80::diCre* was used as recipient strain [7,34].

**Figure 3.**
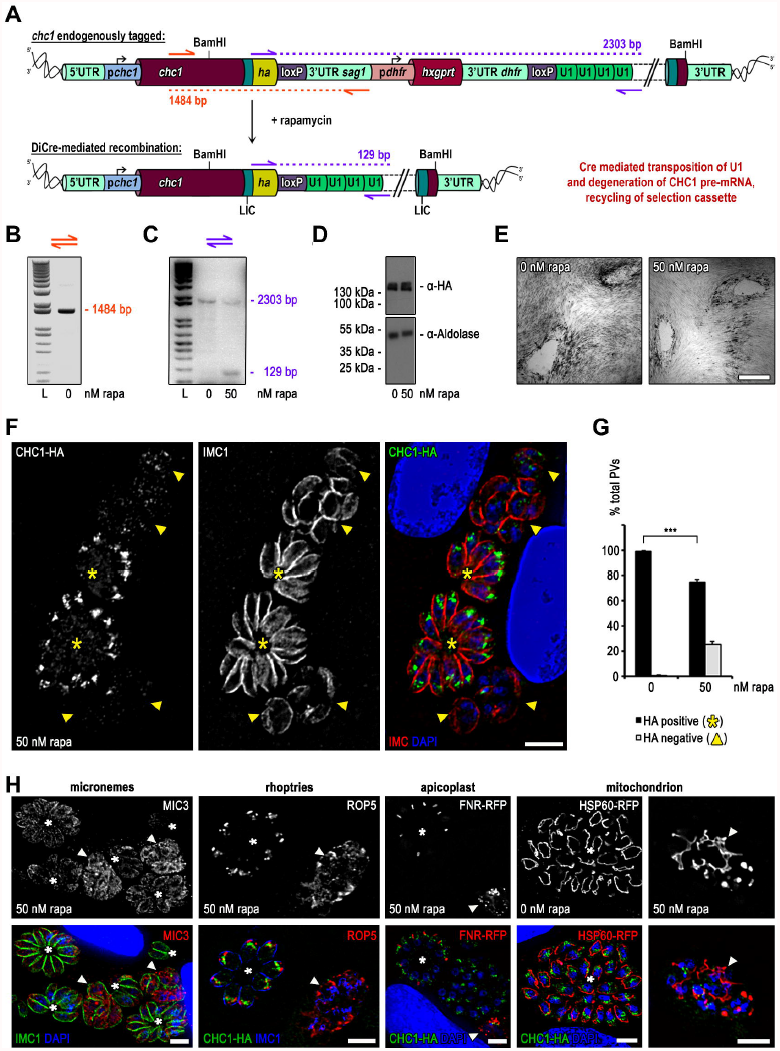
Targeted silencing of *chc1* in *ku80::diCre T. gondii* parasites. (A) Schematics of the genomic locus of *chc1* after single homologous integration of the BamHI linearised endogenous *ha* tagging and U1 gene silencing construct. In absence of rapamycin the four U1 sites are separated from the STOP codon of the inserted tag by a floxed spacer consisting of the 3’ UTR of *sag1* and the *hxgprt* selection cassette. Addition of rapamycin induces Cre recombinase activity. The floxed spacer is excised by site specific recombination and the U1 sites placed adjacent to the STOP codon promoting silencing of *chc1.* (B) Analytical PCR on genomic DNA of clonal CHC1-HA-U1 parasites using oligos indicated as orange arrows in (A). The obtained band of 1484 bp confirms construct integration. (C) Confirmation of Cre/loxP site specific recombination by analytical PCR of genomic DNA. Genomic DNA has been extracted from clonal CHC-HA-U1 parasites grown for 24 h in absence and presence of 50 nM rapamycin. Binding sites of primers used for analysis are indicated with purple arrows in (A). The presence of a specific 2303 bp fragment in absence of rapamycin reflects the genomic composition before Cre/loxP site specific recombination. An additional specific 129 bp fragment in presence of rapamycin verifies successful Cre/loxP site specific recombination. L, ladder. (D) Immunoblot analysis of clonal CHC1-HA-U1 parasites cultured for 24 h with or without 50 nM rapamycin. Membrane probed with anti-HA and anti-Aldolase as loading control. Intensities of detected bands show no difference (E) Giemsa stain. Growth analysis over a 7 days period in absence and presence of 50 nM rapamycin shows no difference in plaque formation. Scale bar represents 500 μm. (F) Immunofluorescence analysis of clonal CHC1-HA-U1 parasites grown for 24 h in presence of 50 nM rapamycin. Green, HA; red, IMC1; blue, Dapi; scale bar represents 10 μM. Endogenous HA-tagged CHC1 localises apical to the nucleus to the trans-Golgi network (yellow asterisks, positive vacuoles). In parasites with silenced *chc1* no specific signal for HA is detectable (yellow arrow heads, HA negative vacuoles). In comparison to HA positive parasites, HA negative parasites show abnormal IMC formation. (G) Quantification of *chc1* silencing efficiency. The graph shows the percentage of HA positive vacuoles (yellow asterisks in (F)) and negative vacuoles (yellow arrow heads in F) determined by examination of 200 vacuoles per condition based on immunofluorescence analyses of clonal CHC1-HA-U1 parasites grown for 24 h in absence or presence of 50 nM rapamycin. Values are means ±SD (n=9). Under rapamycin conditions a significant *chc1* downregulation of 25.4 ± 2.1 was obtained (***, p<0.001, unpaired two-tailed Student’s T-test). (H) Confirmation of specific CHC1 depletion phenotype by immunofluorescence analysis. CHC1-HA-U1 parasites were cultured for 24 h with or without rapamycin. The fate of the inner membrane complex (IMC), micronemes, rhopties, apicoplast and mitochondrion under rapamycin conditions were visualised by using anti-IMC1, anti-MIC3, and anti-ROP5 antibodies and transient transfection of FNR-RFP and HSP60-RFP respectively. In contrast to normal vacuoles (asterisks) silenced vacuoles (arrow heads) show abnormal IMC formation with MIC3 retention in the ER and mislocalisation of ROP5. FNR mislocalises and HSP60 manifests collapsed interlaced mitochondria. Scale bars represent 10 μm.

Previously we demonstrated the consequences of functional interference with CHC1 function using a dominant negative approach [33] and found that clathrin is required for biogenesis of the secretory organelles (micronemes and rhoptries), apicoplast and mitochondria division. We performed a series of immunofluorescence assays on the CHC1-HA-floxU1 parasites after induction with 50 nM rapamycin for 24 hours and confirmed this phenotype (Figure 3F and 3H), demonstrating that U1 mediated silencing is a powerful method to specifically study the function of a GOI. However, a drawback of this approach was the low excision rate. We therefore decided to pursue several alternative strategies to improve excision and to allow a broader application of this single vector strategy.

### A generic knockdown vector that can be used in all strains of *T. gondii*

While the above strategy leads to recycling of the selectable marker, it still requires a recipient strain that expresses both components of the DiCre-system. We reasoned that it should be possible to place the DiCre-expression cassette within the floxed DNA-fragment (Figure 4A). After endogenous tagging of a GOI with an appropriately-designed vector a HA-FLAG tagged version of the respective protein will be expressed. Induction of DiCre will lead to the excision of the selectable marker, the DiCre-expression cassette and positioning of the four U1 recognition sequences downstream of the STOP codon of the GOI (Figure 4A). As a GOI for this strategy we selected the *T. gondii* homolog of yeast VPS26 (TGME49_263500), a putative component of the retromer. Correct integration of the input vector was confirmed by PCR and rapamycin dependent excision of the floxed DNA fragment was confirmed 24 hours after induction with 50 nM rapamycin (Figure 4B and 4C). However, similar to knockdown of CHC1 we were unable to detect significant differences in protein levels at 24 h, 48 h or 72 h after induction with rapamycin (Figure 4B,C and data not shown). Growth analysis did not show obvious differences in plaque formation. Overall, these results indicate a rather low efficiency of recombination. Unexpectedly, we also identified some background excision of the floxed DNA in absence of rapamycin (Figure 4C). Using IFA analysis, we confirmed that VPS26 localises close to the nucleus in Golgi region (Figure 4F), most probably to endosomal-like compartments, similar to TgVPS10 (Sortilin; [35]). Knockdown of TgVPS26 was readily detected in ∼12 % of the induced population 48 h after induction with rapamycin (Figure 4G).

**Figure 4.**
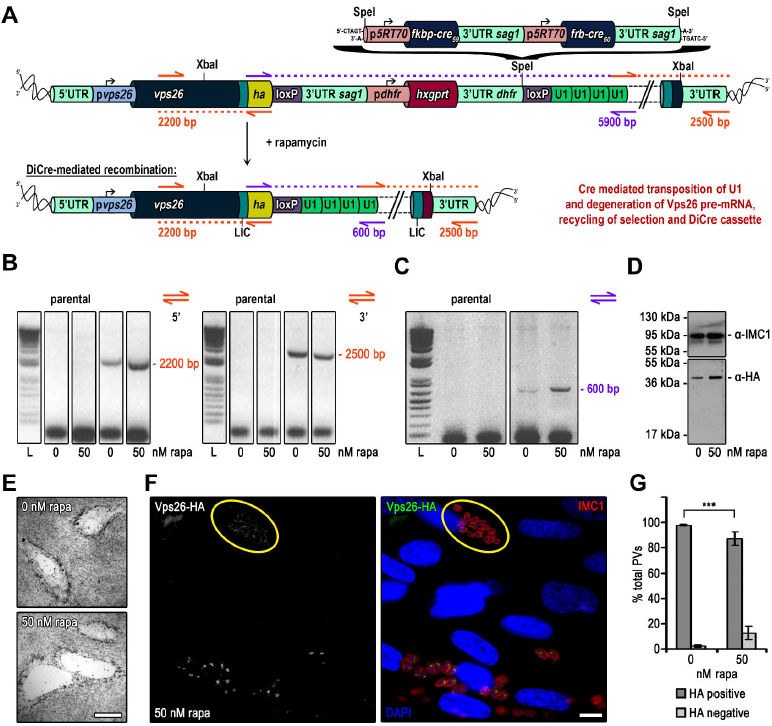
Targeted silencing of *vps26* in *Δku80*-DHFR *T. gondii* parasites. (A) Schematics of the genomic locus of *vps26* after single homologous integration of the XbaI linearised endogenous *ha* tagging and U1 gene silencing construct in absence and presence of rapamycin. A codon optimised DiCre cassette is cloned into the SpeI restriction site between the *hxgprt* selection cassette and the second loxP site. (B) and (C) Analytical PCR on genomic DNA extracted from clonal Vps26-HA-U1 parasites and the parental line Δ*ku80-*DHFR grown for 24 h in absence and presence of 50 nM rapamycin. (B) Construct integration was confirmed by using oligos indicated as orange arrows in (A). Theoretical fragment sizes of 2200 bp and 2500 bp for 5’ and 3’ integration were amplified respectively in Vps26-HA-U1 parasites independent of rapamycin. (C) Confirmation of Cre/loxP site specific recombination. Binding sites of primers used for analysis are indicated with purple arrows in (A) with the theoretical fragment sizes. Even though it was impossible to amplify the very large spacer (5900 bp) in absence of rapamycin the PCR product of 600 bp in presence of rapamycin confirms Cre/loxP site specific recombination. L, ladder. (D) Immunoblot analysis of clonal Vps26-HA-U1 parasites cultured for 48 h with or without 50 nM rapamycin. Membrane probed with anti-HA and anti-IMC1 as loading control. In presence of rapamycin no downregulation of Vps26-HA expression was observed. (E) Giemsa stain. Growth analysis over a 7 days period in absence and presence of 50 nM rapamycin shows no difference in plaque formation. Scale bar represents 500 μm. (F) Immunofluorescence analysis of clonal Vps26-HA-U1 parasites grown for 48 h in presence of 50 nM rapamycin. Green, HA; red, IMC1; blue, DAPI; scale bar represents 10 μM. Endogenous HA-tagged Vps26 localises apical to the nucleus in Golgi region. Parasites with silenced *vps26* are encircled in yellow. (G) Quantification of *vps26* silencing efficiency. The graph shows the percentage of HA positive and negative vacuoles determined by examination of 100 vacuoles per condition based on immunofluorescence analyses of clonal Vps26-HA-U1 parasites grown for 48 h in absence or presence of 50 nM rapamycin. Values are means ±SD (n=3). Under rapamycin conditions a *vps26* downregulation of 12.6 ± 5.32 was obtained (***, p<0.001, Mann-Whitney test).

In summary, while this strategy might be worthwhile to use in different *T. gondii* strains, where no recipient expressing DiCre is available, the induction rate is rather inefficient and therefore within this study we focused on the establishment of new recipient strains, with higher induction rates of DiCre, based on RH-DiCre (7).

### Generation of novel DiCre recipient strains with a high efficiency of rapamycin-dependent recombination

Deletion of *ku80* results in enhanced homologous recombination [8,9] and therefore we developed a strategy to delete the *ku80* locus in the highly efficient RH DiCre-DHFR strain (Figure 5A, [7]) to generate novel strains that would be both highly efficient in homologous recombination and in rapamycin-dependent DiCre recombinase activity. We first replaced the *ku80* coding region with *hxpgrt* (using a cassette in which *hxpgrt* was flanked by 5’ and 3’ UTR of *ku80*) via a homologous double crossover (Figure 5B) as confirmed using genomic PCR (Figure 5E and 5E’). As a second step we transfected *T.gondii* DiCre *ku80::hxgprt* with a construct consisting of the 5’ and 3’ UTR of *ku80* to delete *hxgprt* from the *ku80* locus to generate a “cleaned-up” strain DiCreΔ*ku80* (Figure 5C) or with a construct consisting of 5’UTR *ku80* KRed_flox_YFP 3’UTR *ku80* to generate an indicator strain, where Cre mediated recombination results in a switch from KRed to YFP expression (Figure 5D and 5F). These events were selected using a negative selection reagent, 6-Thioxanthine, for loss of *hxgprt* and were verified by PCR analysis of genomic DNA (Figure 5E’’). We successfully isolated the novel strains DiCreΔ*ku80* and DiCre-*ku80::KRed_flox_YFP*. To assess that the DiCre efficiency had not diminished with the additional genetic manipulations of the RH-DiCre strain, we compared the efficiency of rapamycin induction by transiently transfecting the novel strains DiCre *ku80::hxgprt* and DiCreΔ*ku80* with a reporter plasmid p5RT70-*KRed_flox_-YFP* as previously described [7]. We found the efficiency of rapamycin-induced DiCre recombinase to be comparable to the parental strain (Figure 5G). Similarly, induction of the DiCre-*ku80::KRed_flox_YFP* parasites with rapamycin results in an efficient switch (∼97%) from red to green (Figure 5G). Finally, we compared the expression levels of FRB-Cre59, one of the DiCre subunits, in the different recipient strains and confirmed that expression levels correlate with the efficiency of recombination. Compared to our standard *Ku80::diCre* expression levels are clearly increased in strains based on RH DiCre (Figure 5H, see also [7]).

**Figure 5.**
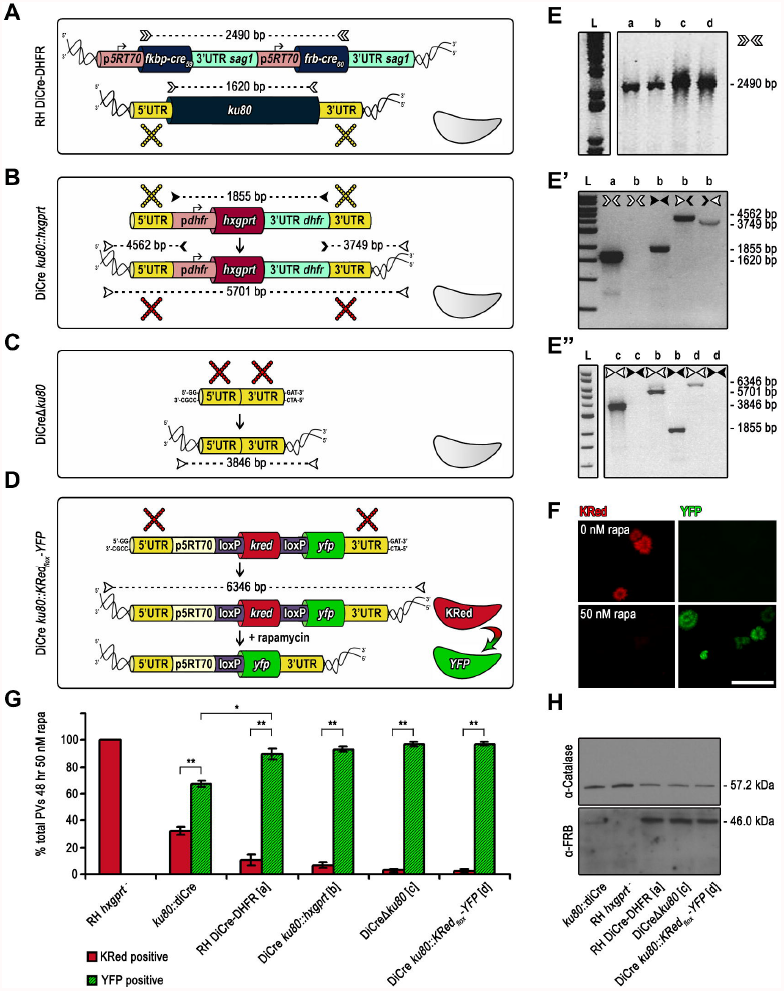
Generation of optimised DiCre expressing recipient and reporter strains DiCre*Δku80* and DiCre *ku80::KRed_flox_-YFP*. (A) Schematics of parental line RH DiCre-DHFR. In this strain the DiCre cassette is randomly integrated into the genome and not targeted into the *ku80* gene locus. (B) Schematics of DiCre *ku80::hxgprt* generation. The *ku80* gene is replaced with the *hxgprt* selection cassette by double homologous recombination and positively selected for HXGPRT using mycophenolic acid and xanthine. Yellow crosses indicate homologous regions in (A) and (B) between which crossovers take place. (C) Schematics of DiCre*Δku80* generation. The *hxgprt* selection cassette is removed by double homologous recombination with an empty knock out construct and negatively selected for HXGPRT using 6-thioxanthine. Red crosses indicate homologous regions in (B) and (C) between which crossovers occur. (C) Schematics of DiCre *ku80::KRed_flox_-YFP* generation. The *hxgprt* selection cassette is replaced with the reporter cassette [7] by double homologous and negatively selected for HXGPRT using 6-thioxanthine. Red crosses indicate homologous regions in (B) and (D) between which crossovers occur. In absence of rapamycin the constitutive p5RT70 promoter drives Killer Red (KRed) expression. Upon addition of rapamycin the floxed open reading frame of *kred* is excised by Cre/loxP site specific recombination and replaced by *yfp*. The shift from red fluorescent to green fluorescent parasites can be used as a measure of Cre recombinase activity which allows discrimination of single parasites. (E-E’’) Analytical PCRs on genomic DNA extracted from the indicated parasite strains. Oligonucelotides used are indicated as arrowheads next to the gel or on top of the lanes. Oligonucleotide binding sites are indicated with the same symbols in (A-D). Predicted PCR product sizes are displayed on black dashed lines between the respective forward and reverse oligonucleotides. Diagnostic PCR for the DiCre cassette was positive in all strains (E). Diagnostic PCRs with different oligonucleotide combinations show successful replacement of the *ku80* locus with the *hxgprt* selection cassette in the DiCre *ku80::hxgprt* strain (E’ and E’’). Diagnostic PCRs confirm cleanup of the *hxgprt* selection cassette in the *ku80* locus of DiCreΔ*ku80* strain and replacement with the reporter cassette in the DiCre *ku80::KRed_fiox_-YFP* strain (E’’). (F) Flourescence analysis of DiCre *ku80::KRed_flox_-YFP* parasites in presence and absence of 50 nM rapamycin for 48 h. Upon rapamycin induction parasite fluorescence shifts from red to green. Scale bar represents 50 μm. (G) Quantification of Cre/loxP site specific recombination efficiency of the DiCre strains. Except for DiCre *ku80::KRed_flox_-YFP* all strains were transiently transfected with the reporter cassette. The graph shows fluorescence analyses of parasites grown for 48 h in presence of 50 nM of rapamycin, the percentage of KRed positive and YFP positive vacuoles were determined by examination of 200 vacuoles in three fields of view. Values are means ±SD (n=4). *, P-Value is <0.0001 *ku80::diCre* vs RH DiCre-DHFR/ DiCre*Δku80*/ DiCre *ku80::KRed_flox_-YFP* (indicated exemplary for the first pair only), **, P-Value is <0.00001 in a 2-tailed Student’s t-test switch from red to green. (H) Immunoblot analysis of indicated parasite strains cultured for 48 h with or without 50 nM rapamycin. Membrane probed with anti-FRB and anti-Catalase antibodies. The latter was used as loading control. FRB-Cre59 subunits show greater abundance in strains with randomly integrated DiCre cassette in comparison to the *ku80::diCre* strainwith a single DiCre cassette in the *ku80* locus.

### Generation of conditional KO mutants for the dynamin related protein C (DrpC)

Next we explored whether the new DiCre recipient strains can be used for efficient U1 dependent silencing. We have previously identified three dynamin related proteins in *T. gondii*, DrpA, DrpB and DrpC [36]. While DrpA has been demonstrated to be required for apicoplast replication [25], DrpB is required for biogenesis of micronemes and rhoptries [36]. In contrast no information is yet available about the highly divergent DrpC and previous attempts to generate GTPase truncated, dominant negative mutants by applying the ddFKBP-system were not successful (Pieperhoff, unpublished data). Therefore we generated an endogenous *drpC* tagging vector analogous to *chc1* described above (Figure 6A) and established endogenously tagged parasite strains in *ku80::diCre* (7), DiCreΔ*ku80* (Figure 5C) and DiCre-*ku80::KRed_flox_YFP* (Figure 5D). Next analytical PCRs were performed on each resulting parasite following treatment with 50 nM rapamycin or mock treatment. In each case correct endogenous tagging was confirmed (Figure 6B-B’’ and 6C-C’’). In the case of *ku80::DiCre*, we observed only inefficient recombination and both the excised and non-excised loci were readily detected in the induced population (Figure 6B), corresponding well to the situation observed for CHC1. In good agreement with this result, no difference in levels of DrpC-HA-FLAG protein were observed in this strain and IFA analysis confirmed that only ∼10% of all parasites showed a knockdown for *drpC* (Figure 6C, 6D and 6E). Consequently, no differences in plaque formation were observed in this knockdown strain (Figure 6F).

**Figure 6.**
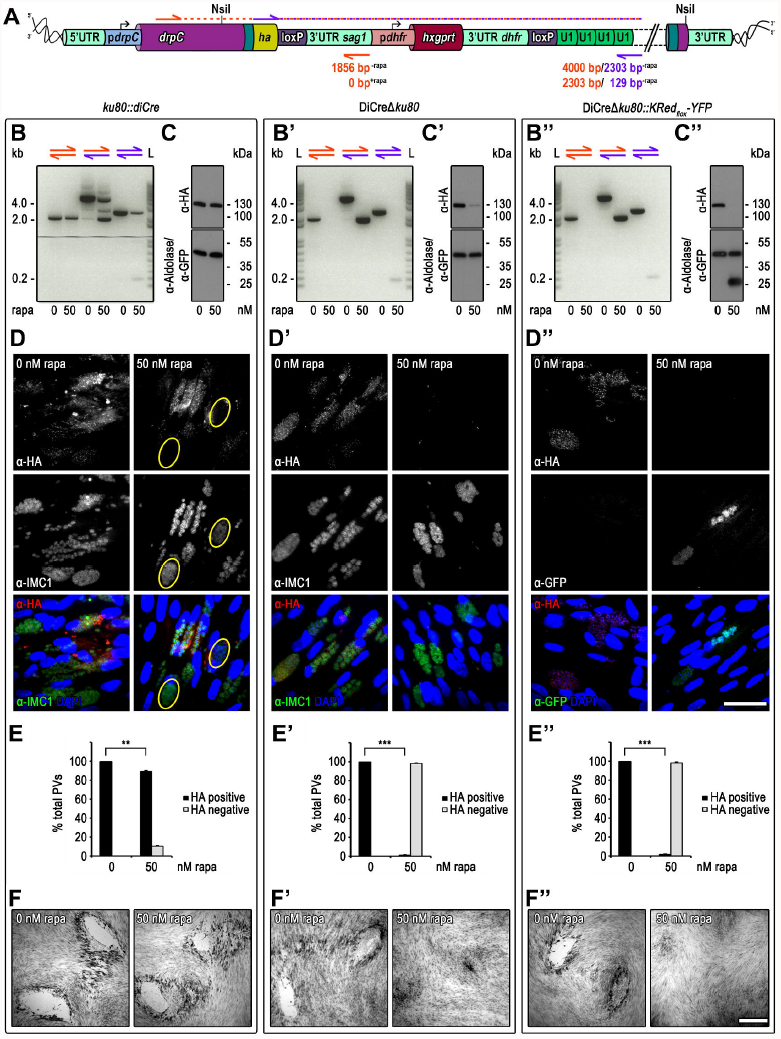
Targeted silencing of *drpC* in *ku80::diCre,* DiCre*Δku80*, and DiCre *ku80::KRed_flox_-YFP* parasites. (A) Schematic of the genomic locus of *drpC* after single homologous integration of the NsiI linearised endogenous *ha* tagging and U1 gene silencing construct in absence of rapamycin. Oligos used for analytical PCRs are illustrated with orange and purple arrows. Theoretical product sizes dependent on rapamycin are indicated below antisense oligos in the colour of the respective sense oligo. Stable construct integration into *ku80::diCre* ([7], B-F), DiCre*Δku80* (B’-F’), and DiCre *ku80::KRed_nox_-YFP* (B’’-F’’). (B-B’’) Analytical integration PCR (orange and mixed oligo combination) and Cre/loxP site specific recombination PCR (all oligo combinations) on genomic DNA extracted from clonal parasite lines after 48 h incubation with or without rapamycin. Resulting PCR product sizes reveal correct integration into all recipient strains, but only complete Cre/loxP site specific recombination in the new recipient strains (B’ and B’’). L, ladder. (C-C’’) Immunoblot analysis of total parasite lysates of clonal lines cultured for 48 h with or without 50 nM rapamycin. Membrane probed with anti-HA, anti-Aldolase, and anti-GFP antibodies. Aldolase was used as loading control. Only new recipient strains show clear rapamycin dependent DrpC-HA expression (C’ and C’’). As expected GFP expression is only detectable in presence of rapamycin in the reporter recipient strain (C’’). (D-D’’) Immunofluorescence analysis of clonal parasites after 48 h incubation with or without 50 nM rapamycin. (D-D’) PFA fixation; green, IMC1; red, DrpC-HA; blue, DAPI (D’’) Methanol fixation to quench KRed and GFP flourescence. GFP is visualised by probing with anti-GFP antibody. Scale bar represents 100 μM. A dotted staining pattern for DrpC-HA was observed in all three recipient strains (D-D’). DiCre*Δku80* (D’) and DiCre *ku80::KRed_flox_-YFP* (D’’) parasites show clearly rapamycin dependent *drpC* silencing. In contrast, in *ku80::diCre* parasites only in a few *drpC* silenced vacuoles (encirceled in yellow) were observed (D). (E-E’’) Quantification of *drpC* silencing efficiency. Graphs show the percentage of HA positive vacuoles and negative vacuoles determined by examination of 200 vacuoles per condition based on immunofluorescence analyses as shown in (D-D’’). Values are means ±SD (n=3). In contrast to a partial *drpC* downregulation of 10.5 ± 0.8 (**, p<0.01, unpaired two-tailed Student’s T-test) in *ku80::diCre* parasites (E), *drpC* was almost completely silenced in DiCreΔ*ku80* (E’; 98.5 ± 0.4; ***, p<0.001, unpaired two-tailed Student’s T-test) and DiCre *ku80::KRed_*flox*_-YFP* (E’’; 98.3 ± 0.6; ***, p<0.001, unpaired two-tailed Student’s T-test) parasites under rapamycin conditions.

In strong contrast, knockdown of *drpC* in both DiCreΔ*ku80* and DiCre-*ku80::KRed_flox_YFP* was highly efficient. Indeed, PCR analysis suggested complete removal of the floxed DNA sequence in the induced population and only a faint band for DrpC-HA-FLAG was still detectable in immunoblot analysis 24 hours after induction (Figure 6B’, 6C’, 6B’’ and 6C’’). Furthermore in immunofluorescence analysis we could hardly detect any parasites that showed a clear signal for DrpC-HA (Figure 6D’ and 6D’’) and we confirmed that none of the YFP positive parasites were also positive for HA, demonstrating that DiCre mediated recombination of the reporter plasmid is a good indicator for recombination in a different locus (Figure 6D’’). Recombination rate in both cases was close to 100% (Figure 6E’ and 6E’’) and no plaque formation was observed in the presence of rapamycin (Figure 6F’ and 6f’’). These results confirm the high efficiency of recombination that can be obtained with these recipient strains. In summary, U1 dependent silencing can be efficiently combined with DiCre mediated recombination to achieve generation of GOI knockdown mutants in a fast and efficient manner.

### The U1 mediated knockdown strategy is not functional in *P. falciparum*

Encouraged by our results in *Toxoplasma,* we attempted to adapt the U1 mRNA destabilisation strategy to another apicomplexan parasite, the human malaria pathogen *Plasmodium falciparum*. As a target GOI in this organism we selected *pfsub1* (PlasmodDB ID PF3D7_0507500), which encodes a subtilisin like serine protease involved in parasite egress from the infected host erythrocyte. Based on previous failed attempts to disrupt this gene in *P. falciparum* [28], as well as conditional knockout studies of the orthologous gene in the rodent malaria species *P. berghei* [37,38], *pfsub1* is thought to be an essential gene in parasite blood-stages. For modification of the *pfsub1* gene, we created a plasmid construct designed to integrate by single crossover homologous integration into the genomic locus, introducing a C-terminal HA3 epitope tag as well as a floxed heterologous 3’ UTR (the PbDT 3’ UTR) downstream of the *pfsub1* ORF. A sequence comprising an array of 10 repeated U1 recognition sequences was placed immediately downstream of the floxed PbDT3’ UTR (Figure 7A). The construct was transfected into the previously-described *P. falciparum* 1G5DiCre recipient clone [6], which constitutively expresses DiCre from a genomic locus. Rapamycin induced DiCre mediated recombination was predicted to excise the PbDT 3’ UTR, concomitantly translocating the 10 U1 recognition sequences to a position directly adjacent to the *pfsub1* STOP codon. We expected that loss of the 3’ UTR along with addition of the U1 sequences would lead to mRNA destabilisation and result in PfSUB1 protein knockdown. Correct integration of the construct at the *pfsub1* locus in the 1G5DiCre *P. falciparum* parasites and epitope tagging of the PfSUB1 gene product was confirmed by diagnostic PCR (not shown) and by IFA using the anti 3HA mAb 3F10 (Figure 7B). Integrant parasites were cloned by limiting dilution and two independent clones (D3 and B11) selected for detailed analysis. Complete excision of the floxed sequence was observed within 44 h of rapamycin treatment of the clones (Figure 7C), confirming the previously-reported high efficiency of DiCre mediated excision in *P. falciparum*. Nucleotide sequencing of the excision-specific PCR product confirmed deletion of the PbDT 3’ UTR and resulting translocation of the 10 U1 recognition sequences to just downstream of the *pfsub1* coding sequence.

**Figure 7.**
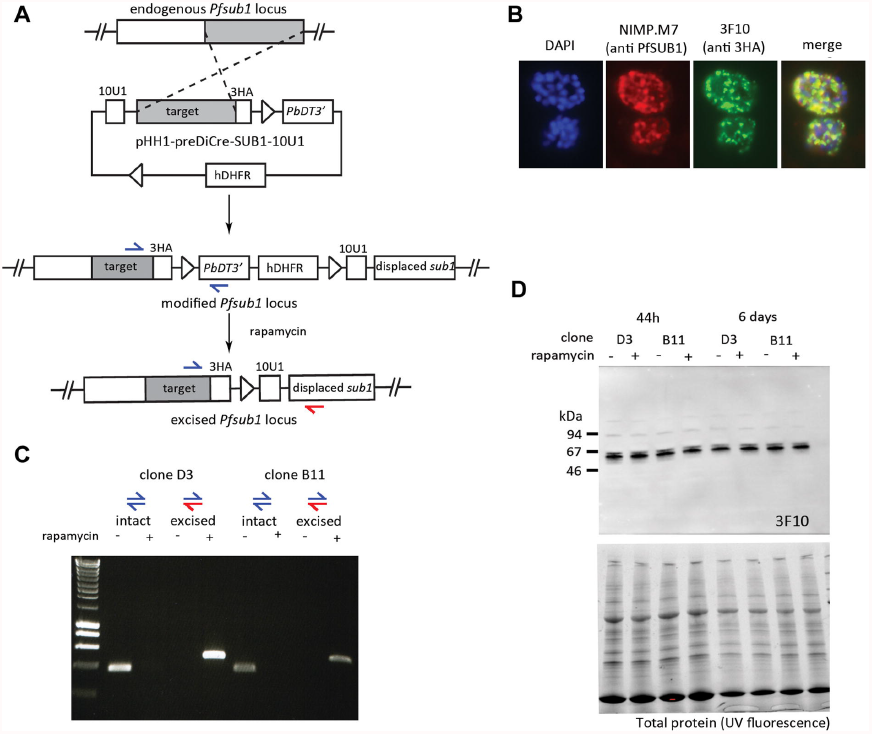
The U1 mediated knockdown strategy is not functional in *P. falciparum.* (A) Schematic showing the strategy used to replace the endogenous *pfsub1* 3’ UTR with a floxed heterologous *PbDT* 3’ UTR with a downstream sequence comprising 10 U1 recognition sequences. The single cross-over homologous recombination event also results in the fusion of a HA3 epitope tag to PfSUB1. The predicted results of DiCre mediated recombination induced by treatment with rapamycin are shown. The positions of hybridization of PCR primers designed to specifically amplify sequences from the intact and excised locus are indicated with red and blue arrows. (B) IFA of mature schizonts of integrant clone D3, probed with mAb NIMP.M7 (anti PfSUB1) and mAb 3F10 (anti HA3). (C) Efficient DiCre-mediated recombination between the integrated loxP sites, as shown by diagnostic PCR analysis of parasite genomic DNA extracted 44 h following rapamycin treatment of two independent parasite clones D3 and B11. The blue and red arrows correspond to primer pairs marked in (A). (D) Upper panel: Western blot analysis with mAb 3F10 shows no down-regulation of PfSUB1 protein expression protein 44 h or 6 days after rapamycin treatment. Lower panel: loading controls, showing total protein SDS PAGE gels detected by UV fluorescence on the BioRad Chemidoc™ XRS system prior to Western transfer.

To test the effects of the recombinase mediated genomic alterations on PfSUB1 expression, mature schizonts from rapamycin-treated and control cultures were analysed by IFA (not shown) and Western blot 44 h following rapamycin treatment (Figure 7D). Surprisingly, no discernible effects on PfSUB1 protein levels were observed in either clone (Figure 7D). To determine whether this might be due to the expression of *pfsub1* mRNA early in that erythrocytic cycle (i.e. prior to rapamycin-treatment) we maintained the treated parasites for an additional two erythrocytic cycles, analysing schizonts once again six days following rapamycin treatment. The parasite cultures replicated at the same rate, and examination by Western blot again found no detectable differences in PfSUB1 expression levels between the control and the rapamycin-treated parasites (Figure 7D) . This result suggested that despite its efficacy in *Toxoplasma*, U1 mediated knockdown is not functional in *P. falciparum*.

In further work, a similar strategy was taken to attempt conditional knockdown of two additional *P. falciparum* genes, subtilisin like protease 2 (*pfsub2*: PlasmodDB PF3D7_1136900) and merozoite surface protein1 (*msp1*: PlasmoDB). In both cases, despite similarly efficient DiCre mediated excision of the inserted 3’ UTR and translocation of the 10 U1 recognition sequences adjacent to the STOP codon, no effects on protein expression was observed, strongly suggesting key differences in regulation of splicing in these two apicomplexan genera.

### Discussion

In this study we successfully established a new strategy that allows the generation of conditional knockdown *T.gondii* parasites using a one vector approach. We transferred knowledge from studies on pre-mRNA processing in other eukaryotic systems that demonstrates that interference with exon-intron definition by placing U1 recognition sequences in the terminal exon [31] results in mRNA degradation. Indeed a technology based on U1 mediated gene silencing has been developed [32], but has the drawback that multiple genetic manipulations are required.

Here we provide evidence that the basic mechanisms of pre-mRNA splicing must be conserved between *T. gondii* and other eukaryotes, since positioning of the highly conserved U1 recognition sequences into the terminal exon (or close to the STOP codon) of *T. gondii* genes results in gene silencing.

This allowed us to combine the advantages of DiCre mediated recombination [7,15] with U1 mediated gene silencing by conditionally placing U1-recognition sequences next to the STOP codon of genes of interest (GOI). In order to improve this system we also developed new *T.gondii* DiCre recipient strains that have a low background excision in the absence of the inducer rapamycin, but more than 90% excision efficiency when rapamycin is added, for as little as 4 hours, to the parasite culture.

In order to test the efficiency and specificity of this novel technology, we generated conditional knockdown parasites for three *T.gondii* GOIs, *chc1, vps26* and *drpC.* Importantly, comparison of the phenotype of *chc1-*knockdown parasites with a previously described dominant negative mutant for CHC1 [33] did not show any obvious differences, indicating that the knockdown effect is specific for *chc1* gene function. We also attempted a one vector approach that can be applied in all strains of *T. gondii* since the expression cassette for DiCre is placed into the tagging vector and is removed along with the selectable marker. However, although successful targeting of *vps26* could be achieved, we only obtained a population with ∼12% knockdown parasites for this essential gene. This is likely due to poor expression levels of DiCre and therefore the successful application of this strategy most likely depends on the genomic location of the GOI. Indeed, when we compared different DiCre-expressing strains, we found a strong correlation of DiCre expression levels and efficiency of recombination, indicating that for successful application of DiCre strong expression of the individual subunits (Cre59 and Cre60) should be ensured. Therefore it is our suggestion to limit this approach for the manipulation of parasite strains where no transgenic DiCre-expressing recipients are available. However, it is important to mention that even knockdown efficiencies of less than 10% can be useful to characterise a phenotype, since the respective parasites can be easily identified within a mixed population.

Another drawback of the technology presented here is the fact that integration of loxP sequences within the 3’-UTR appears to directly influence gene expression, resulting in reduced protein levels, as observed in case of mycGFP (see Figure 2). We suspect this may be one of the reasons, why several attempts to tag a GOI with the U1-vector failed and the success rate at this point is approximately 20% (not shown). However, this limitation can likely be overcome by using different 3’UTR sequences in future versions of this technology. Unfortunately we failed to directly adapt the U1 degradation system to use in *P. falciparum.* At this point we are not sure why this strategy should not work in this closely related parasite, given that U1 recognition sequences are completely conserved between *T. gondii* and *P. falciparum.* Future investigations and optimisations are required to solve this puzzle. Importantly, this technology can easily be applied in a high-throughput approach and can be combined with the recently adapted CRISPR/Cas9 system in *T. gondii* [39].

## Material and Methods

### *T. gondii* parasite lines, maintenance and transfections

*T. gondii* RHΔ*hxgprt*, RH BloxP DiCre [7], *ku80::diCre* [7], and Δ*ku80*-DHFR [8] tachyzoites were maintained by serial passage in human foreskin fibroblast (HFF) monolayers cultured in Dulbecco’s modified Eagle’s medium (DMEM) supplemented with 10% fetal bovine serum (FBS), 2 mM glutamine, and 25 μM gentamycin at 37°C and 5% CO_2_ in a humidified incubator. Transfections were carried out by electroporation as previously described [16] using approximately 10^7^ freshly egressed or mechanically released parasites of the respective parasite strain.

### Generation of *T. gondii* expression constructs and stable *T. gondii* parasite lines

#### GFP-U1 reporter parasite lines

For the generation of the plasmids p5RT70mycGFP-U1-HX and p5RT70mycGFP-mutU1-HX complementary oligo pairs (U1-for/U1-rev and mutU1-for/mutU1-rew) with protruding overhangs were annealed after prior phosphorylation and ligated into the PstI and PacI restriction sites of p5RT70mycGFPDD-HX [4]. Hereby the c-terminal ddFKBP tag was removed and a STOP codon for the myc-GFP cDNA was inserted followed downstream by two U1 (CAGGTAAGTT) or two mutated U1 (mutU1, CATGGAACTT) recognition sites respectively. The p5RT70mycGFP-floxU1-HX plasmid was generated by amplification of a floxed 3’ untranslated region (UTR) using oligo pair floxU1-for/floxU1-rev with p5RT70mycGFP-U1-HX as template. In addition to the flanking loxP sites, the resultant PCR product possessed a STOP codon for the myc-GFP cDNA at the 5’ end and two U1 recognition sites at the 3’ end. After digestion with the restriction enzymes PstI and NotI it was ligated into the respective restriction sites of p5RT70mycGFP-U1-HX. 30 μg of the respective plasmid were linearised with SacI and co-transfected with 15 μg of the SacI linearised pBSSK^+^SAG1/BLE/SAG1 plasmid [17] into RH BloxP DiCre parasites [7]. The resultant transfectants were selected for clonal lines expressing mycGFP-U1, mycGFP-mutU1 and mycGFP-floxU1 by phleomycin selection using a combination of high-dose (50μg/ml) extracellular treatment and low-dose intracellular treatment (5 μg/ml) as previously described [18] and subsequent cloned by limiting dilution.

### CHC1-HA-floxU1 parasite line

For C-terminal HA-FLAG epitope endogenous tagging and U1 mediated knockdown of the *chc1* gene an EcoRV flanked cassette with a ligation-independent cloning (LIC) cassette [8] 5’ to the start of the HA-FLAG cDNA and a loxP flanked 3’ UTR of SAG1 and HXGPRT selection cassette placed adjacent to the STOP codon of the HA-FLAG cDNA followed by four U1 recognition sites was synthesised and inserted into pUC57-Simple by GenScript USA Inc. (Piscataway, NJ 08854, USA). This plasmid is referred to as pLIC-HA-FLAG-(3’UTR_SAG1_-pDHFR-HXGPRT-5’UTR_DHFR_)_flox_-4xU1. The 3′ flank of the *chc1* gene upstream of the STOP condon was amplified by polymerase chain reaction (PCR) from *T. gondii* RH Δ*hxgprt* strain genomic DNA using oligo pair CHC1-LIC-for/CHC1-LIC-rev and inserted into pLIC-HA-FLAG-(3’UTR_SAG1_-pDHFR-HXGPRT-5’UTR_DHFR_)_flox_-4xU1 by ligation-independent cloning strategy [8]. 15 μg of the resultant CHC1-HA-FLAG-(3’UTR_SAG1_-pDHFR-HXGPRT-5’UTR_DHFR_)_flox_ plasmid were linearised by BamHI within the region of homology for efficient homologous recombination and were transfected into *ku80::diCre* parasites [7]. The resultant transfectants were selected for clonal lines expressing CHC1-HA in presence of 25 μg/ml mycophenolic acid and 40 μg/ml xanthine as previously described [19] and subsequent cloned by limiting dilution. Specific integration and DiCre mediated site specific recombination were confirmed by analytical PCR on genomic DNA using oligo pair CHC1-integr-for/ pG152-integr-rev and pG152-ssr-for/pG152-ssr-rev respectively.

### DrpC-HA-floxU1 parasite lines

For generation of the DrpC-HA-FLAG-(3’UTR_SAG1_-pDHFR-HXGPRT-5’UTR_DHFR_)_flox_-4xU1 plasmid the *drpC* gene 3′ flank was PCR amplified on *T. gondii* RHΔ*hxgprt* strain genomic DNA with oligo pair DrpC-LIC-for/DrpC-LIC-rev. The resultant PCR product was inserted into pLIC-HA-FLAG-(3’UTR_SAG1_-pDHFR-HXGPRT-5’UTR_DHFR_)_flox_-4xU1 by LIC [8]. 10 μg NsiI linearised plasmid each were transfected into *ku80::diCre* [7], DiCre *ku80*, and DiCre Δ*ku80::KRed_flox_-*YFP parasites. Resulting transfectants were selected for HXGPRT [19] and cloned by limiting dilution. Specific integration and DiCre mediated site specific recombination were confirmed by analytical PCR on genomic DNA using oligo pairs DrpC-integr-for/pG152-integr-rev, pG152-ssr-for/pG152-ssr-rev, and DrpC-integr-for/pG152-ssr-rev.

### Vps26-HA-floxU1-DiCre parasite line

A codon optimised DiCre cassette was synthesised by GenScript USA Inc. (Piscataway, NJ 08854, USA) and inserted into the SpeI restriction site of pLIC-HA-FLAG-(3’UTR_SAG1_-pDHFR-HXGPRT-5’UTR_DHFR_)_flox_-4xU1 to generate the plasmid pLIC-HA-FLAG-(3’UTR_SAG1_-pDHFR-HXGPRT-5’UTR_DHFR_-DiCre)_fl__ox_-4xU1. The 3’ flank of *vps26* was PCR amplified on *T. gondii* RHΔ*hxgprt* strain genomic DNA with oligo Vps26-LIC-for and Vps26-LIC-rev and inserted into pLIC-H A-FL AG-(3’ UTR_SAG1_-pDHFR-HXGPRT -5’UTR_DHFR_-DiCre)_flox_-4xU1 by LIC. 15 μg of the XbaI linearised plasmid were stable transfected into Δ*ku80-*DHFR [8] tachyzoites. Resulting transfectants were selected for HXGPRT [19] and cloned by limiting dilution. Specific integration and DiCre mediated site specific recombination were confirmed by analytical PCR on genomic DNA using oligo pairs V ps26-integr-5’-for/V ps26-integr-5’-rev, Vps26-integr-3’-for/V ps26-integr-3’-rev and pG152-ssr-for/Vps26-ssr-rev.

### DiCre optimised recipient and reporter parasite lines

The parental line RH DiCre-DHFR was generated by stable transfection of 60 μg p5RT70DiCre-DHFR into RHΔ*hxgprt* tachyzoites as described in Andenmatten et al. 2013 (7) previously. Transfectants were selected for dihydrofolate reductase (DHFR) with 1 μM pyrimethamine as described previously [20] and subsequent cloned by limiting dilution. Integration was confirmed by diagnostic PCR on genomic DNA with oligo combination DiCre-for/DiCre-rev.

To generate DiCre *ku80::hxgprt* parasites the *ku80* gene was replaced with the *hxgprt* cassette by double homologous recombination. Briefly, 60 μg of 5’Ku80-pDHFR-HXGPRT-3’Ku80 PCR product (8) was transfected into RH DiCre-DHFR parasites. Transfectants were selected for HXGPRT [19] and cloned by limiting dilution. *ku80* replacement by the *hxgprt* cassette was confirmed by analytical PCR on genomic DNA with oligo combinations Ku80-for/Ku80-rev, HXGPRT-for/HXGPRT-rev, Ku80-5’-for/HXGPRT-5’-integr-rev, and HXGPRT-3’-integr-for/Ku80-3’-rev.

DiCre *ku80* was generated by double homologous recombination with the knock out construct 5’Ku80-3’Ku80. Plasmid 5’Ku80-DiCre-3’ku80 (7) was modified using SpeI. 60 μg of the religated plasmid were digested with SacII and EcoRV prior to transfection into DiCre *ku80::hxgprt* parasites. Transfectants were cultured in presence of 340μg/ml 6-thioxanthine to select for loss of *hxgprt* [21] and clones were isolated by limiting dilution.

Genomic DNA of individual clones was screened with oligonucleotide combinations Ku80-for/Ku80-rev and HXGPRT-for/HXGPRT-rev.

Cre/loxP site specific recombination efficiency of isolated RH DiCre-DHFR, DiCre *ku80::hxgprt,* and DiCreΔ*ku80* clones was monitored and confirmed by transient transfection with the DiCre reporter plasmid p5RT70-*KRed_flox_-YFP* [7].

Furthermore the stable DiCre reporter line DiCreΔ*ku80::KRed_lox_-YFP* was generated. Briefly, the *hxgprt* cassette was replaced with the reporter cassette 5’Ku80-p5RT70-*KRed_flox_-YFP*-3’Ku80 by double homologous recombination. 60 μg of the reporter cassette were transfected into DiCre *ku80::hxgprt* parasites which were selected for loss of *hxgprt* as above[21]. Clones were isolated by limiting dilution. Genomic DNA of individual clones was screened with oligonucleotide combinations Ku80-for/Ku80-rev and HXGPRT-for/HXGPRT-rev.

### Fluorescence and immunofluorescence microscopy of *T. gondii* parasites

For microscopic analysis confluent monolayers of HFF cells seeded onto 13 mm diameter glass cover slips were infected with the respective parasite line and incubated with or without 50 nM rapamycin at normal growth conditions.

For fluorescence analysis cells were fixed after 48 h with 4% w/v paraformaldehyde in phosphate buffered saline (PBS) for 20 min at room temperature, rinsed with water and mounted onto microscope slides with DAPI-Fluoromount-G (SouthernBiotech). Samples were examined with a Carl Zeiss Axioskope 2 MOT Plus inverted epifluorescence microscope fitted with a 10x objective lens (Plan-APO-CHROMAT, NA 0.45) and equipped with a Hamamatsu Photonics Orca-ER CCD digital camera. Images were acquired with Improvision OpenLab 5.0 software and processed with Adobe Photoshop CS4 Extended software.

For immunofluorescence analysis cells were fixed after 24 h or 48 h either with 4% w/v paraformaldehyde in PBS for 20 min at room temperature or with absolute methanol pre­cooled to -20°C for 20 min at -20°C. Fixed cells were permeabilised with 0.2% Triton X-100 in PBS for 20 min and blocked with 3% w/v bovine serum albumin (BSA) in permeabilisation buffer for 20 min. Cells were incubated for 1 h at room temperature with anti-HA-Tag (6E2) mouse monoclonal antibody (Cell Signaling Technology, Inc.) diluted 1:100, anti-HA-Tag (3F10) rat monoclonal antibody (Roche) diluted 1:500, anti-TgIMC1 rabbit antiserum [22] diluted 1:1000-1500, anti-TgMIC3 (T42F3) mouse monoclonal antibody [23] diluted 1:100, anti-TgNtRop5 (T53E2) mouse monoclonal antibody [24] diluted 1:1000, and anti-GFP mouse monoclonal antibody (Roche) diluted 1:500 in blocking buffer. Cells were washed three times with PBS and incubated for 1 h at room temperature with Alexa Fluor 488-, Alexa-594-and/or Alexa Fluor 350-conjugated secondary antibodies (Molecular Probes, Invitrogene) diluted 1:3000 (or 1:1000 for DrpC-HA detection) in blocking buffer. Cover slips were washed with PBS, rinsed with water and mounted onto microscope slides with DAPI-Fluoromount-G (SouthernBiotech). For co-localisation studies with apicoplast and mitochondrion parasites were transiently transfected with 50 μg of TgHSP60-RFP-CAT [25] or TgFNR-RFP-CAT (21) plasmid DNA prior to the experiment. For overview images samples were examined with a Carl Zeiss Axioskope 2 MOT Plus inverted epifluorescence microscope fitted with a 40x (Plan-APO-CHROMAT, NA 0.45) or 100x oil immersion lens (Plan-APO-CHROMAT, NA 1.4) objective lens. Images were acquired with a Hamamatsu Photonics Orca-ER CCD digital camera using Improvision OpenLab 5.0 software and processed with Adobe Photoshop CS4 Extended software. For high resolution images z-stacks of 0.2 μm increments were collected on a DeltaVision Core epifluorescence microscope (Applied Precision, GE) fitted with a 100x oil immersion lens (UPlanSApo, NA 1.40) and equipped with a Photometrics CoolSNAP HQ2 CCD digital camera using Applied Precision softWoRx Suite 2.0 software. Deconvolution was performed with Applied Precision softWoRx Suite 2.0 software and deconvolved images were further processed with ImageJ 1.44 and Adobe Photoshop CS4 Extended software.

### Immunoblot analysis of total *T. gondii* parasite lysates

For immunoblot analysis parasites were maintained in absence or presence of 50 nM rapamycin for 24 or 48 h. Freshly egressed or mechanically released parasites were harvested, washed once in ice cold PBS, and resuspended in Laemmli sample buffer containing 2% SDS and 100 mM DTT. Total parasite lysates were boiled for 5 min at 95°C and spun down for 5 min at 21.100 x g at room temperature. Supernatant volumes appropriate to 1, 2.5 or 5x 10^6^ parasites were separated on 6 or 13% sodium dodecyl sulphate polyacrylamide gels or 4-20% gradient Mini-Protean TGX precast polyacrylamide gels (BioRad) at 200 V, and transferred onto nitrocellulose membranes. Membranes were blocked with 3% w/v dried skimmed milk and 0.2% Tween 20 in PBS and probed with anti-HA-Tag (6E2) mouse monoclonal antibody (Cell Signaling Technology, Inc.) diluted 1:500, anti-HA-Tag (3F10) rat monoclonal antibody (Roche) diluted 1:2000, anti-c-Myc (9E10) mouse monoclonal antibody (Santa Cruz Biotechnology, Inc.) diluted 1:500, anti-TgIMC1 rabbit antiserum [22] diluted 1:10000 or 1:20000, anti-TgCatalase (84) rabbit antiserum [26] diluted 1:3000, and anti-TgAldolase (WU1614) polyclonal antibody [27] diluted 1:10000 or 1:20000, anti-GFP mouse monoclonal antibody (Roche) diluted 1:1000, anti-ddFKBP12 rabbit polyclonal antibody (Thermo Scientific) diluted 1:500, anti-mTor (human FRB Domain) rabbit polyclonal antibody (Enzo Life Science) diluted 1:250 in blocking buffer. Primary antibodies were detected with Peroxidase-conjugated AffiniPure Goat Anti-Rabbit IgG (H+L), Donkey Anti-Mouse IgG (H+L) (Jackson ImmunoResearch Laboratories, Inc.) and/or Anti-Rat (Pierce antibodies) secondary antibodies respectively diluted 1:50.000 in blocking buffer containing 0. 4% Tween 20. For detection immunoblots were treated with Amersham ECL Plus or Prime Western Blotting Detection Reagents and exposed to Kodak General Purpose Blue Medical X-Ray Films.

### Plaque assay

For growth and viability analysis of the respective parasite lines HFF monolayers were grown in six-well cell culture plates, infected with 50 or 500 parasites per well, and cultured for 7 days in absence or presence of 50 nM rapamycin under normal growth conditions. Cells were fixed with absolute methanol pre-cooled to -20°C for 10 min at room temperature, stained for 30 min with Giemsa which was washed off intensely with PBS. Samples were examined with a Carl Zeiss Axiovert 40 CFL inverted epifluorescence microscope fitted with a 4x objective lens (ACHROPLAN, NA 0.1) and equipped with AxioCam Icc1 CCD digital camera (Zeiss). Images were captured with AxioVision 4.8 (Zeiss) software and processed with Adobe Photoshop CS4 Extended software.

### Statistics

Per condition 100 to 200 parasitophorous vacuoles (PVs) were examined with a Carl Zeiss Axiovert 40 CFL inverted epifluorescence microscope fitted with a 100x oil immersion lens (A-PLAN, NA 1.25). Percentages of mean values of at least three independent experiments ± SD were determined and plotted in a bar graph. P-values were calculated using unpaired two­tailed Student’s t-test unless stated otherwise.

### Maintenance and synchronisation of *P. falciparum*

Asexual blood stages of *P. falciparum* clone 1G5DiCre (ref 6) were maintained and synchronized using standard protocols [28] in RPMI 1640 medium containing Albumax (Invitrogen). Mature schizonts were purified from highly synchronous cultures using Percoll (GE Healthcare) as described previously [29].

Generation of construct pHH1-preDiCre-SUB1-10U1 and transfection into *P. falciparum* Plasmid construct pHH1-preDicre-10U1 was generated using vector pHH1_SERA5del3preDiCre (6) as a template. A synthetic linker region of 249 bp was synthesised in pUC57 (GenScript) to contain 10 adjacent U1 sequences, a multiple cloning site and a LIC region between *Avr*II and *Xho*I restriction sites. Digestion of pHH1_SERA5del3preDiCre using *Spe*I and *Xho*I and the synthetic region with *Avr*II and *Xho*I allowed for the ligation of the synthetic fragment, while enabling the recycling of both the *Avr*II and *Spe*I sites in the multiple cloning site to create the pHH1-preDiCre-10U1 intermediate vector. Plasmid pHH1-preDicre-10U1 therefore contains a floxed PbDT 3’ UTR followed by 10 U1 recognition sequences and the human dihydrofolate reductase (*hdhfr*) cassette which confers resistance to the antifolate drug WR99210. Plasmid construct pHH1-preDiCre-SUB1-10U1 was produced by excising a *pfsub1* targeting region fused to a 3HA tag from plasmid pPfSUB1HA3 [28] using HpaI and XhoI restriction sites, and ligating it into pHH1-preDicre-10U1 digested with the same enzymes. The resulting plasmid was confirmed to contain the *pfsub1* targeting insert by restriction digest and by sequencing. 10 μg of plasmid DNA was used to transfect schizonts and integrants selected by drug cycling as described previously [6]. Integration of the plasmid construct was confirmed by PCR analysis and by IFA using the anti-HA3 mAb 3F10. Integrant parasites were cloned by limiting dilution and two integrant, B11 and D3, were selected for further examination.

### Immunofluorescence assay (IFA) and Western blot

IFA was performed on Percoll purified schizonts as described [6] using the anti PfSUB1 mAb NIMP.M7 and the anti-HA mAb 3F10 (Roche). For Western blot analysis, purified schizonts were solubilised in SDS sample buffer and fractionated on a 4-15% Biorad Mini PROTEAN^®^ Stain-free ™ gradient SDS polyacrylamide gel. The resolved proteins were visualised by UV fluorescence on the BioRad Chemidoc™ XRS system prior to electrophoretic transfer onto nitrocellulose membrane. For Western analysis the blot was probed as described previously [30] using mAb 3F10. Chemiluminescent signal was visualised using the BioRad Chemidoc™ XRS system.

### Rapamycin treatment and PCR analysis of *P. falciparum* clones

Highly synchronous ring-stage cultures of integrant parasite clones D3 and B11 were treated with 100 nM rapamycin for 4 h. The rapamycin was washed off and the parasites returned to culture for 44 h (when they reached mature schizont stage) or for up to 6 days. Genomic DNA was prepared from the rapamycin treated and the control (mock-treated) parasites as described previously [6]. The intact modified *pfsub1* locus was amplified by PCR using primers endo_SUB1_FOR_end and PbDT3’R1. The excised locus (following DiCre-mediated recombination) was amplified using primers endo_SUB1_FOR_end and endo_SUB1_REV 1.

**Table 1.**
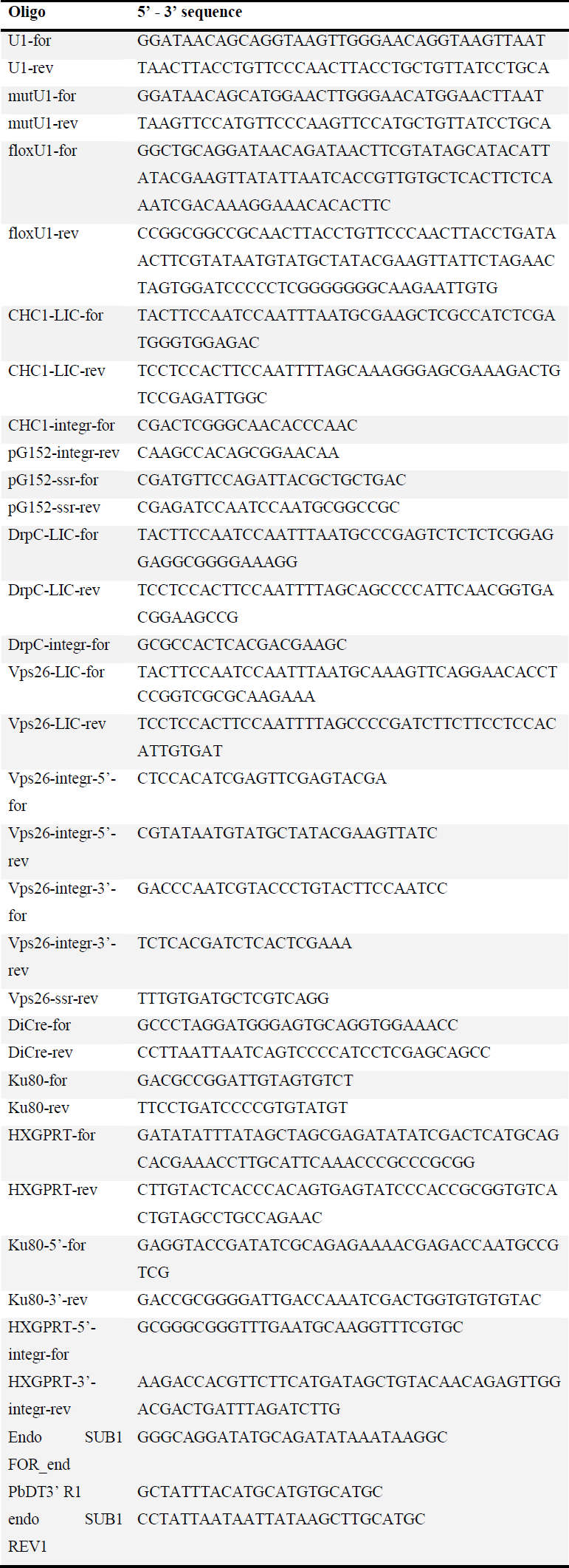
Oligonucleotides used for cloning of *T. gondii* and *P. falciparum* constructs, confirmation of construct integration and site-specific recombination. LIC, ligation independent cloning; for, forward; rev, reverse; mut, mutated; ssr, site specific recombination; integr, integration

**Table 2.**
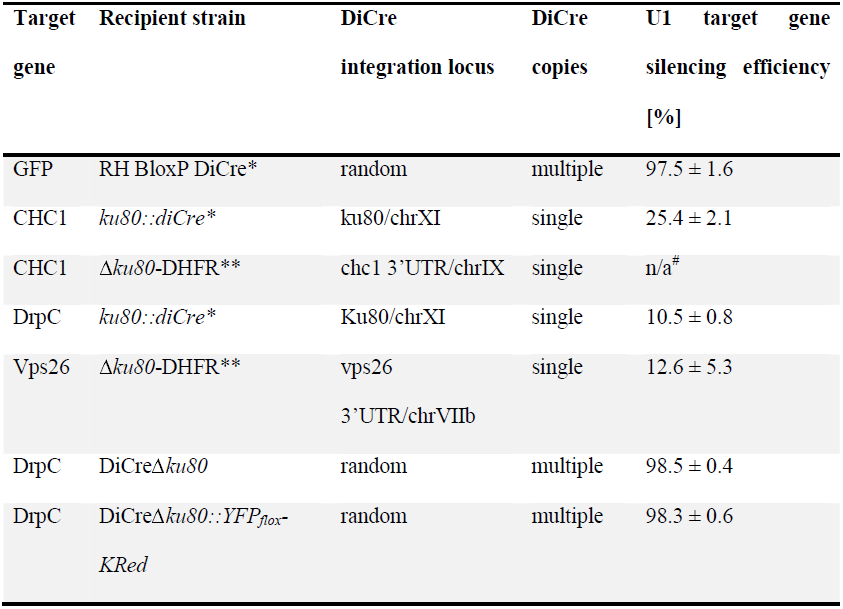
Correlation of U1 gene silencing efficiency to genomic locus and copy numbers of DiCre. Chr, Chromosome; *, [40]; **, [8]; *#*, Stable integration was not achieved in five independent transfections.

## Acknowledgement

We would like to thank Samuel I. Gunderson (Rutgers University, USA) for valuable input during the initial phase of this project.

